# Spotted fever *Rickettsia* and relapsing fever *Borrelia* in rodents from southern India

**DOI:** 10.64898/2026.02.14.705948

**Authors:** B.R. Ansil, Tejas Pawar, Pallab Majee, Rridhi Kapila, Tikily Libang, Uma Ramakrishnan

**Affiliations:** National Centre for Biological Sciences, Tata Institute of Fundamental Research, Bengaluru, Karnataka, India; School of Biological Sciences, University of Oklahoma, Norman, OK, USA; Department of Microbiology and Cell Biology, Indian Institute of Science, Bengaluru, Karnataka, India

**Keywords:** zoonoses, vector-borne pathogens, coinfection, *Leptospira*, *Orientia*, *Coxiella*

## Abstract

Bacterial zoonoses constitute a substantial fraction of emerging infectious diseases. In this study, we investigated the presence of pathogenic bacterial genera—*Rickettsia, Borrelia, Orientia, Leptospira*, and *Coxiella*—in rodents from southern India. We detected low circulation of *Rickettsia* (7.26%), *Borrelia* (6.45%), and *Leptospira* (0.8%), whereas *Orientia* and *Coxiella* were not detected in the rodents sampled. Notably, we observed contrasting patterns of tissue association, with rickettsiae detected exclusively in pooled organ tissues and borreliae detected only in blood, suggesting the influence of pathogen biology in detection probabilities. Our phylogenetic analyses further revealed spotted fever group rickettsiae and relapsing fever group borreliae in both synanthropic and forest-associated rats, highlighting a potential transmission risk to people and livestock in the region. By revealing the circulation of zoonotic bacteria in new host species, this study underscores the need for systematic surveillance of wildlife to better characterize bacterial diversity and its public health implications.

## 1. Introduction

Zoonotic pathogens maintained in wildlife reservoirs represent a persistent and growing threat to public health and global economies (1). Rodents, the most species-rich mammalian order comprising rats, mice, and squirrels, are among the most important wildlife reservoirs to disproportionately host several zoonotic pathogens (2). This distinctive reservoir role of these taxa arises from their large, dynamic populations and exceptional adaptability to human-modified environments, which together facilitate the maintenance and transmission of pathogens (3). Although we do not see large rodent-associated bacterial pandemics, emerging infections such as rickettsioses, Lyme disease, and leptospirosis are continuing to cause significant public health burden, especially in tropical countries (4,5).

Anthropogenic disturbances, including deforestation, agricultural expansion, and urbanization, have profoundly transformed natural habitats and reshaped rodent community composition and abundance, often favoring generalist and synanthropic species (5,6). These ecological shifts can fundamentally affect host–vector–pathogen interactions by modifying host densities, contact rates, and vector assemblages, thereby increasing spillover opportunities to humans and other sympatric animals (5). Understanding how zoonotic bacteria are distributed across rodent diversity and habitats is therefore critical for identifying zoonotic risk and predicting how environmental change may influence disease emergence. Despite this clear link between rodent ecology and zoonotic emergence, a significant gap exists in the characterization of rodent-associated bacterial pathogens and their enzootic cycles. This paucity of systematic surveillance, particularly in tropical countries like India, severely constrains our understanding of pathogen diversity, ecology, and transmission pathways.

India harbors high rodent diversity with 110 species spanning 48 genera (7), representing substantial zoonotic reservoir potential. Rodent-associated infections are of particular concern in India due to significant human exposure to rodents and arthropod vectors largely driven by agrarian livelihoods and human–animal interactions (both wildlife and farming associated) in rural and peri-urban landscapes (8). Zoonotic surveillance efforts focused on human populations across India have documented pathogens such as *Rickettsia* spp., *Borrelia* spp., *Orientia tsutsugamushi, Coxiella burnetii, Leptospira* spp., and many more (9–12). Rodents are known reservoirs of these primarily vector-borne bacteria (except *Leptospira* spp.), causing significant public health burden (13). For example, rickettsioses caused by *Rickettsia spp*. and scrub typhus caused by *O. tsutsugamushi* cause serious infection and can be life-threatening if not treated (9). These infections are prevalent in rural India but are often under- or misdiagnosed due to overlapping clinical presentations with other common infections (14). However, systematic investigations of zoonotic bacterial pathogens in India remain limited to a small fraction of the rodent diversity. In particular, host specificity and the extent of pathogen co-circulation within individual hosts have rarely been examined. Previous studies in southern India detected *Staphylococcus aureus* and *Salmonella typhimurium* in synanthropic rodents (16). Additionally, our earlier work in the region documented a high prevalence of *Bartonella* spp. in forest-associated rodents, including both host-specific and zoonotic lineages (15). These findings point to a broader, largely uncharacterized landscape of bacterial pathogen diversity circulating among Indian rodents.

In this study, we contribute towards this knowledge gap by investigating the occurrence of clinically important rodent-associated zoonotic bacterial genera within a forest–plantation mosaic in southern India. Based on previous detections in rodents and documented human infections, we focused on five bacterial genera of public health relevance—*Rickettsia, Borrelia, Orientia, Leptospira*, and *Coxiella*. We examined the prevalence, phylogenetic diversity, and evolutionary placement of these bacterial taxa. We further assessed the influence of host species identity and sex on infection prevalence and coinfection patterns. We expected high prevalence of these bacteria in the focal rodent community, with sex-specific differences in infection prevalence. Additionally, we also expected frequent coinfections, particularly among synanthropic species such as *Rattus rattus*.

## 2. Methods

### 2.1 Rodent sampling

We sampled rodents (rats and mice) in Kadamane, a forest-plantation mosaic in Karnataka, southern India, between January and March 2018 as part of a broader study on small mammal-associated pathogens in the Western Ghats (15,17). Sampling followed a grid-based framework using 100 medium-sized Sherman traps arranged in a 10×10 configuration to cover one hectare grid. Five forest grids, two grassland grids, and one human habitation grid were surveyed, with trapping effort allocated proportionally to each habitat type within the study area (Figure 1A). Captured individuals were humanely euthanized for sample collection (NCBS-IAEC-2016/10-[M]), and species identification was confirmed using a combination of morphological and molecular methods (15). We collected blood as dried blood spots (DBS) on FTA cards and organ tissues (liver, spleen, kidney, lung, and intestine) in RNAlater. Samples were stored at 4°C for about one week before being transferred to −20°C at the National Centre for Biological Sciences (NCBS).

**Figure 1.**
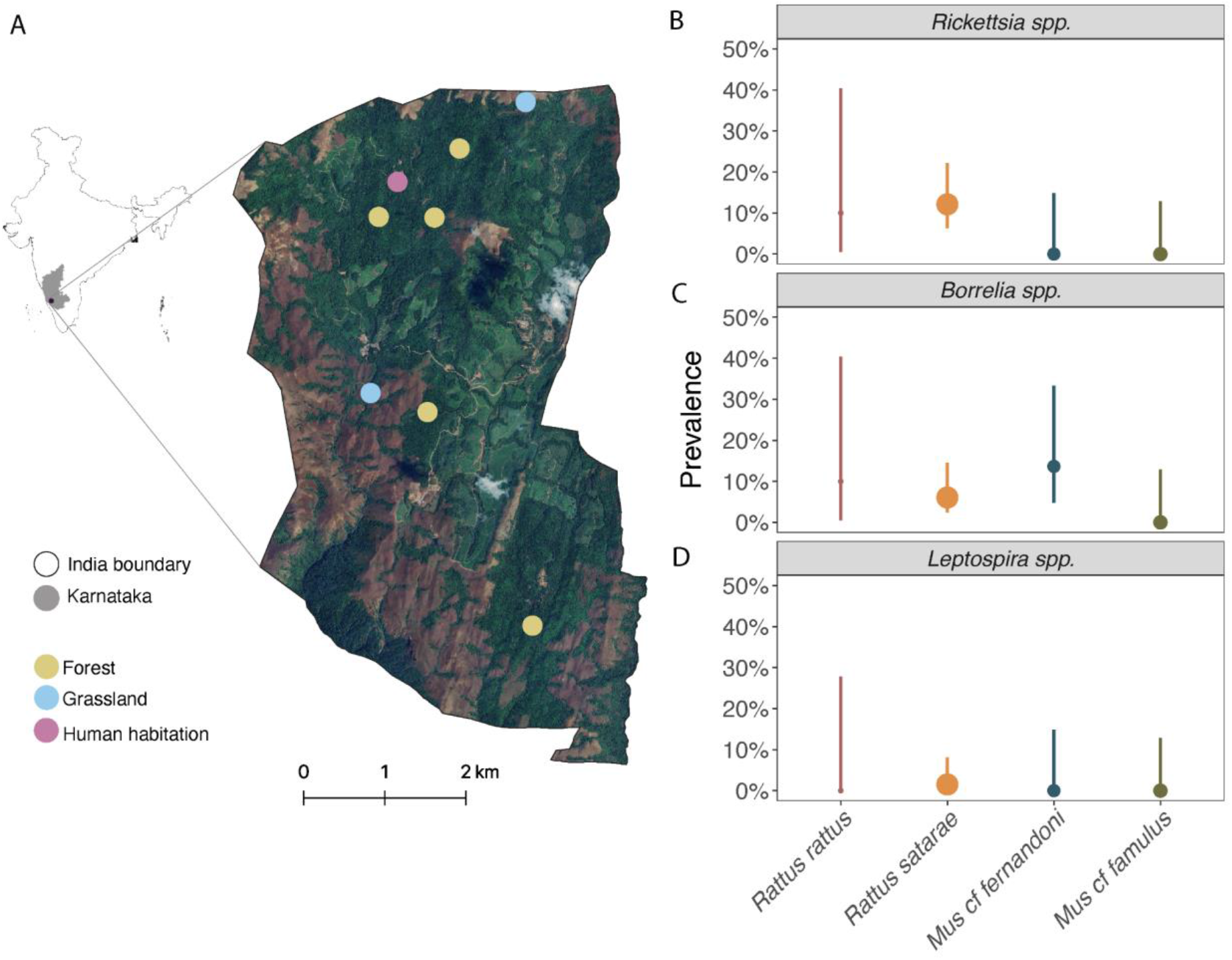
Map of the study area (left) showing sampling points colored by habitat types where small mammals were trapped. Prevalence of *Rickettsia, Borrelia*, and *Leptospira* (right) in tested samples is shown with point estimates and 95% confidence intervals. Point sizes are scaled by sample size.

### 2.2 Molecular diagnostics

Among the seven species captured (17), only four species with adequate sample sizes (n ≥ 10) were included in this study (Table S1), representing *Rattus rattus* (n = 10), *Rattus satarae* (n = 66), *Mus cf. fernandoni* (n = 22), and *Mus cf. famulus* (n = 26). We extracted DNA from DBS using the QIAamp DNA Mini Kit (Qiagen). These extracts were used to screen for *Rickettsia, Borrelia*, and *Orientia* using previously described PCR assays targeting *OmpB* (811 bp), 16S rRNA (650 bp), and 56-kDa protein (371 bp) loci, respectively (18–20). Similarly, DNA from kidney and spleen was extracted using the Quick-DNA/RNA Miniprep Plus Kit (Zymo Research) and screened for *Leptospira* (16S rRNA; 289 bp) and *Coxiella* (IS1111A; 146 bp), respectively (21,22). Additionally, approximately five grams of each organ (liver, spleen, kidney, lung, and intestine) were subsampled, pooled, and homogenized using a mechanical bead beater. DNA was then extracted from these homogenates using the method mentioned above and screened for all five bacterial genera. This pooled-tissue screening approach was included to increase the probability of detection of bacterial agents that might have been missed when screening only specific organs or DBS.

All PCRs were performed in a 25 μL reaction volume containing 12.5 μL of 1× HotStarTaq Master Mix (Qiagen), 1 μL each of 5 μM primers, 6.5 μL of nuclease-free water, and 4 μL of template DNA. *Borrelia* and *Leptospira* assays employed a nested PCR approach, in which the second-round reactions used 1 μL of first-round product and 9.5 μL of nuclease-free water. All reactions were run for 40 cycles, and amplicons were visualized on 1.5% agarose gels stained with GelRed (Biotium Inc.). Primer sequences and PCR conditions are provided in Table S2. PCR results were interpreted with reference to the positive and negative controls included in each batch of samples. Previously sequence confirmed samples were used as positive controls for *Borrelia, Rickettsia, Orientia* and, *Leptospira* whereas extracted DNA from established culture was used as the positive control for *Coxiella*. Amplicons were purified and sequenced at NCBS Sanger sequencing facility. Sequences were then aligned and consensus sequences were generated in Geneious Prime (https://www.geneious.com/). Identity for each sequence was then confirmed using BLAST searches against the NCBI reference database.

### 2.3 Statistical analysis

Individuals were classified as positive if a bacterium was detected in any of their samples. We then estimated the prevalence (proportion of positive individuals in a species) and 95% confidence intervals using the Hmisc package in R (23). We fitted two generalized linear models (GLMs), each with binary responses and mean bias reduction using the brglm2 package in R (24) to assess the effects of species, sex, and their interaction on the probability of *Rickettsia* and *Borrelia* infection. The first model included only additive effects, while the second included the species and sex interaction. The best-supported models were selected after comparing corrected Akaike Information Criterion (AICc), ΔAICc, number of coefficients (*k*) and Akaike weights (*wi*) (Table S3). We then applied a Wald test (25) to evaluate the significance of each predictor on infection status. Since species occurrence was uneven across habitats, (Table S1), we did not include ‘habitat’ as a covariate in our models.

We incorporated previously reported *Bartonella* occurrence data from these rodents (15) into our dataset to examine coinfection within individual rodents. We used pairwise coinfection status (*Rickettsia-Borrelia, Rickettsia-Bartonella*, and *Borrelia-Bartonella*) as binary response variables in three additional GLMs. In addition to species and sex, we also incorporated the occurrence of the non-focal bacteria as an additional predictor. All the GLMs were fitted and evaluated as mentioned above.

### 2.4 Phylogenetic analysis

Consensus sequences from each bacterial genus from this study, along with reference sequences of known species from GenBank, were aligned using MUSCLE in Geneious Prime. All three alignments were subsequently cleaned using Gblocks, best nucleotide substitution models were assessed, and maximum-likelihood phylogenies were constructed with PhyML on the NGPhylogeny.fr platform (26). Each phylogeny was generated with 10,000 bootstrap iterations. Phylogenies were then visualized, annotated, and unique lineages were identified in FigTree v1.4.4 (27). Additionally, we selected representative sequences from each unique phylogenetic lineage and pairwise genetic dissimilarity (p-distance) relative to their closest known species were computed using the ape package (28). Dissimilarity was then converted to similarity (1− dissimilarity) and represented as percentage.

## 3. Results

### 3.1 Bacterial infection in rodents

We detected rickettsial infection in nine of 124 rodents tested (Figure 1B, Table S3). Specifically, only two species of rats were detected with rickettsiae: *R. rattus* (10%; 95 % confidence interval [CI]: 0.51–40.41%) and *R. satarae* (12.12%; 95 % CI: 6.27–22.13%). The detection was solely on pooled tissues and none of the blood samples were positive (Table S3). These two host species showed contrasting habitat associations, with *R. rattus* detected exclusively in human habitations, whereas *R. satarae* was restricted to forest fragments (Table S1). Our top GLM and Wald test revealed no significant association between rickettsial infection and host identity (*χ*^*2*^=3.50, p=0.32) or sex (*χ*^*2*^=0.16, p=0.68). This model outperformed the alternative model that included a species and sex interaction (Table S4).

Borrelial infections were detected in eight individuals belonging to three species (Figure 1C, Table S3): *R. rattus* (10%; 95% CI: 0.51–40.41%), *R. satarae* (6.06%; 95% CI: 2.38–14.57%), and *M. cf. fernandoni* (13.63%; 95% CI: 4.74–33.33%). All detections were from blood samples, and none of the pooled tissue samples tested positive. Similar to *Rickettsia*, our top *Borrelia* GLM and Wald tests indicated no significant association between infection and host identity (χ^2^=3.18, p=0.36) or sex (χ^2^=0.51, p=0.47).

*Leptospira* prevalence was low in the rodents tested (Figure 1D), with only one *R. satarae* individual testing positive (1.51%; 95% CI:0.07–8.09%). Notably, positivity was detected in both kidney and pooled tissue samples (Table S3), likely due to the inclusion of kidney tissue in the pooled samples. No *Orientia* and *Coxiella* were detected in any of the samples tested.

### 3.2. Coinfection in rodents

We observed 1.61% (n=2; 95% CI:0.44–5.69%) *Rickettsia*-*Borrelia* coinfection among the rodents tested (Figure S1). We observed no significant associations between *Rickettsia-Borrelia* coinfection and host species (χ^2^=2.47, p=0.47), sex (χ^2^=0.15, p=0.69), or the presence of *Bartonella* (χ^2^=0.07, p=0.78). Similarly, *Rickettsia-Bartonella* coinfection was detected in 4.03% of individuals (n=5; 95% CI:1.73–9.09%), with no significant associations observed with host species (χ^2^=1.94, p=0.58), sex (χ^2^=0.29, p=0.58), or the presence of *Borrelia* (χ^2^=2.33, p=0.12). *Borrelia-Bartonella* coinfection was detected in a small proportion of individuals (2.41%; n=3; 95% CI:0.82–6.87%) and showed no significant associations with host species (χ^2^=0.99, p=0.80), sex (χ^2^=0.47, p=0.49), or the presence of *Rickettsia* (χ^2^=2.86, p=0.09). Finally, triple coinfection with *Rickettsia, Borrelia*, and *Bartonella* was rare, occurring in a single *R. satarae* individual (0.81%; 95% CI:0.04–4.46%). No individuals were detected with coinfection by all four bacteria (*Rickettsia, Borrelia, Leptospira*, and *Bartonella*).

### 3.3 Genetic diversity of Rickettsia and Borrelia

Our *OmpB* locus-based phylogeny revealed two distinct rickettsial lineages circulating in rats of the genus *Rattus* (Figure 2A). The single sequence obtained from *R. rattus* formed the first lineage (R1) along with *Rickettsia massiliae* (AF123714), a spotted fever group (SFG) *Rickettsia*. Corroborating this placement, both BLAST and *p*-distance estimates showed 100% sequence similarity between R1 and *R. massiliae* (Figure 2B). The remaining eight identical sequences derived from *R. satarae* formed the second lineage (R2) clustering with *R. honei* (AF123724), another SFG *Rickettsia*. The high sequence similarity (99.7%; Figure 2B) between these sequences and *R. honei* suggests that they likely share the same taxonomic identity.

**Figure 2.**
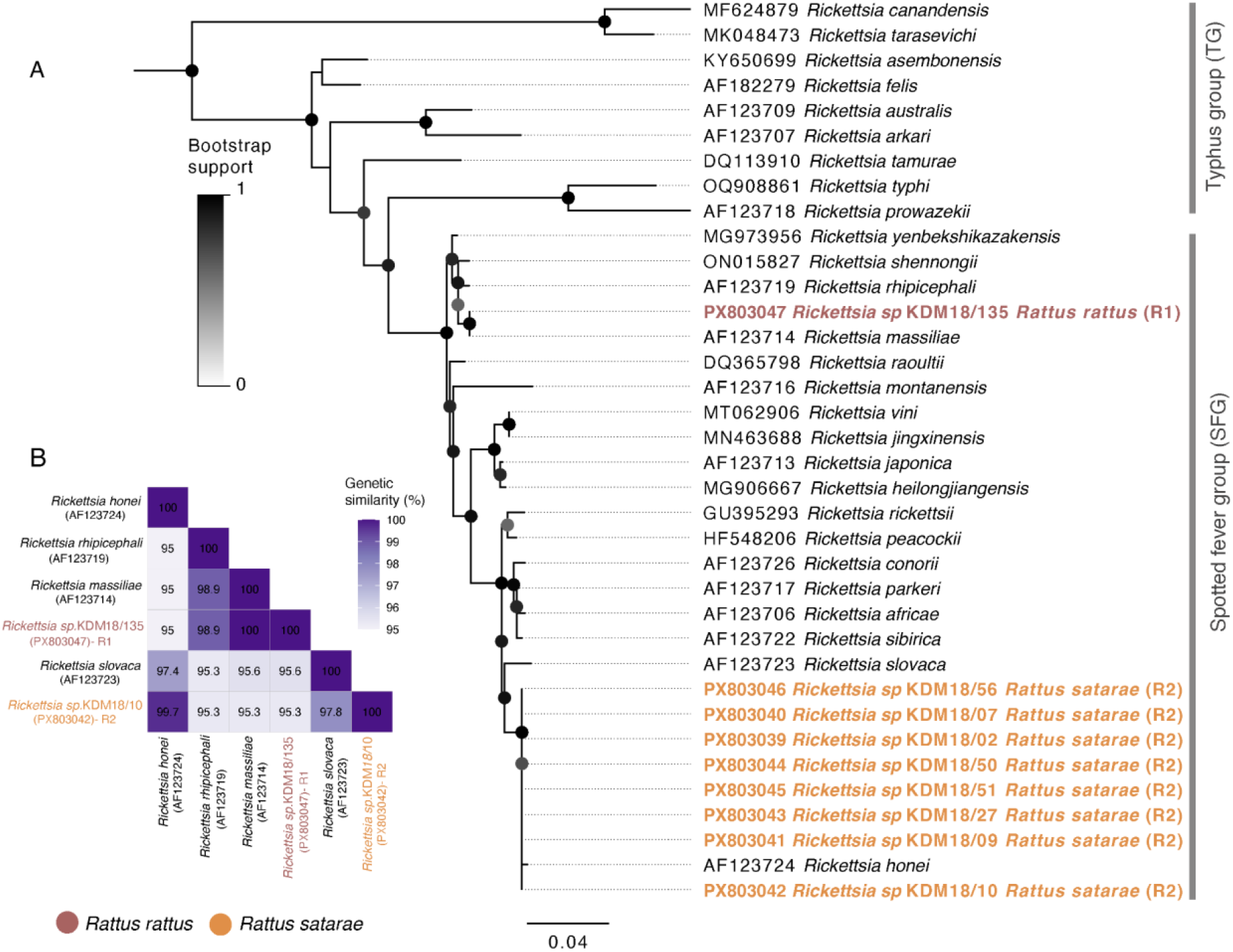
(A) Maximum-likelihood phylogenetic tree of *Rickettsia* spp. based on ompB sequences, constructed using 10,000 bootstrap replicates under the GTR +G substitution model. (B) Genetic similarity of the *Rickettsia* sequence derived from this study in comparison with closely related *Rickettsia* species.

The 16S rRNA phylogeny of *Borrelia* sequences revealed five distinct lineages (Figure 3A) circulating in the host community. Lineages one, two, and three (B1, B2, and B3) formed a unique monophyletic cluster positioned basal to the relapsing fever group (RFG) and Lyme disease group (LDG) *Borrelia*. These three lineages shared 92.0–94.2% sequence similarity with *Borrelia puertoricensis* (NR_181316), whereas, pairwise similarity among B1–B3 was higher, ranging from 94.4 to 95.9% (Figure 3A; Figure 3B). The B1 comprised two identical sequences derived from *M. cf. fernandoni* and was positioned basal to B2 and B3. Lineage B2 consisted of a single divergent sequence from another *M. cf. fernandoni* individual and was basal to B3. Notably, B3 comprised two identical sequences derived from *R. rattus* and *R. satarae*, (Figure 2A), suggesting co-circulation of this lineage in both host species. The fourth lineage (B4) consisted of two identical sequences from *R. satarae* and exhibited high genetic similarity to *B. theileri* (99.6%; LC656242), a RFG *Borrelia*. The fifth lineage (B5) comprised a single sequence from *R. satarae* and formed a sister taxon to *B. turcica* (96.5%; AB111854), occupying an intermediate phylogenetic position between RFG and LDG *Borrelia*.

**Figure 3.**
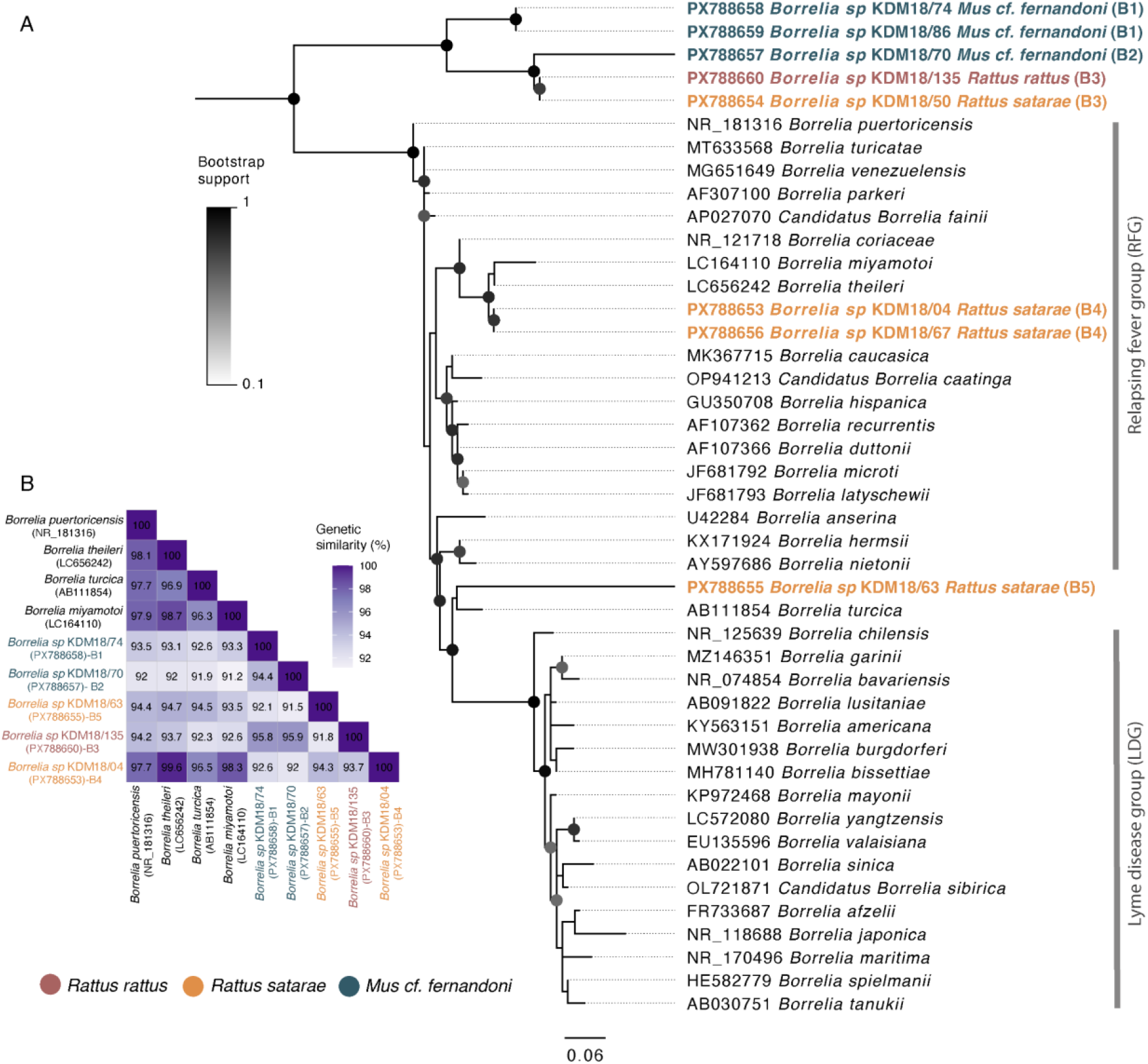
(A) Maximum-likelihood phylogenetic tree of *Borrelia* spp. based on 16S rRNA gene sequences, constructed using 10,000 bootstrap replicates under the GTR + G +I substitution model. (B) Genetic similarity of the *Borrelia* sequence derived from this study in comparison with closely related *Borrelia* species.

### 3.4 Phylogenetic position of Leptospira sequence

16S rRNA sequences analysis of *Leptospira* revealed a distinct phylogenetic placement for the single sequence detected from *R. satarae* (Figure S2A). This sequence was nested within the pathogenic *Leptospira* clade and positioned basal to *Leptospira interrogans* (AY461870) and *L. kirschneri* (NR_043051), while exhibiting the highest genetic similarity (96.4%) to *L. borgpetersenii* (AM050576; Figure S2B), a pathogenic species with multiple serovariants and global distribution.

## 4. Discussion

Characterizing pathogenic diversity in host taxa is fundamental to understanding zoonotic risk. We investigated infection patterns and genetic diversity of five bacterial pathogens (*Rickettsia, Borrelia, Orientia, Leptospira*, and *Coxiella*) in a rodent community sampled across contrasting habitats in southern India. *Rickettsia* and *Borrelia* were detected at low prevalence in rodents inhabiting both human habitation and natural habitats. *Leptospira* infection was rare, detected in a single individual, while *Orientia* and *Coxiella* were not detected in any of the rodents screened. Coinfections involving two bacterial taxa were infrequent, and higher-order coinfections were rare among wild rodents from southern India.

### 4.1. Low but detectable circulation of SFG Rickettsia and RFG Borrelia in rodents

Studies investigating rickettsial and borrelial pathogens in Indian rodents remain limited. Besides, existing works have largely focused on rickettsial agents and have predominantly employed serological methods (e.g., (29). Investigations targeting *Borrelia* in rodents are notably absent in India, despite human cases and detection in ectoparasites (30,31), highlighting a critical gap in characterizing enzootic cycles of these agents. PCR-based studies from other Asian countries report prevalence levels comparable to those observed here, suggesting a broader pattern of low but persistent circulation of rickettsial and borrelial pathogens within rodent assemblages (32,33). However, higher positivity of both *Rickettsia* and *Borrelia* was reported in arthropod vectors, indicating their crucial role in the enzootic cycle and transmission among rodents and other incidental hosts (32,34).

The two *Rickettsia* lineages detected in this study belonged to SFG, and exhibited high sequence similarity to *R. massiliae* and *R. honei*, both of which are widely distributed species known to cause human infections (35,36). Previous clinical studies in India have reported SFG rickettsiae, including *R. massiliae* (but not *R. honei*), from acute encephalitis syndrome patients and rodent-associated ectoparasites (37,38). The high frequency of occurrence of the *R. honei*-like lineage in our dataset (seven of eight detections) suggests that this lineage (R2) could be common but host-specific (*R. satarae*) in South India. Serological studies on Lyme disease in India have primarily implicated *B. burgdorferi* as the causative agent (10,31). Our study, however, identified divergent *Borrelia* lineages, including both potentially novel taxa and members of the RFG, with one lineage (B4) exhibiting high genetic similarity to *B. theileri*, a species known to infect wildlife and livestock (39). These findings suggest that rodents in India may harbor a broader diversity of *Rickettsia* and *Borrelia* than previously recognized.

### 4.2. Contrasting tissue-specific associations and diversity of Rickettsia and Borrelia

We observed contrasting tissue association patterns for *Rickettsia* and *Borrelia*, reflecting fundamental differences in their biology. *Rickettsia* was detected exclusively in pooled tissues and not in blood, probably due to its obligate intracellular lifecycle and the typically transient or low-level bacteremia associated with infection (40). Consequently, rickettsial detection in peripheral blood may require assays with greater sensitivity (40). In contrast, *Borrelia* was detected solely in blood samples, consistent with its extracellular nature and its ability to proliferate in the bloodstream during active infection, a process that facilitates efficient vector-mediated transmission (41). Such differences in tissue-specific association and methodological choices may explain variability in prevalence across studies (42). However, obtaining internal tissue samples is not always logistically feasible and may not be appropriate when hosts of interest are threatened or endangered. In such contexts, targeting minimally invasive tissues (e.g., ear clips) alongside host-associated ectoparasites may provide a practical alternative for detection and characterizing enzootic transmission cycles (43).

The phylogenetic analyses further revealed low genetic diversity in *Rickettsia*, with two SFG lineages (Figure 2A) showing host-specific patterns. Specifically, *R. massiliae* was associated with the synanthropic *R. rattus*, whereas *R. honei* was associated with the forest-dwelling *R. satarae*, suggesting ecological structuring and a potential barrier for cross-species transmission (44). Nevertheless, the limited sample size for *R. rattus* and the reliance on a single genetic locus constrain the strength of this inference. Broader sampling across hosts, vectors, and genetic loci will be necessary to confirm these associations. In contrast to *Rickettsia, Borrelia* exhibited high genetic diversity, with five distinct lineages detected, including multiple lineages co-circulating within *R. satarae* and *M. cf. fernandoni* (Figure 3A). Such patterns of population-level co-circulation are not uncommon in *Borrelia* spp. (45). Moreover, the detection of *Borrelia* in *M. cf. fernandoni*, while the complete absence of *Rickettsia* may reflect differences in host competence towards these agents, or vector specificity, collectively suggests a limited role for mice in maintaining rickettsial infections while potentially contributing substantially to borrelial maintenance in the rodent community.

Notably, despite occupying contrasting habitats, *R. rattus* (in human settlements) and *R. satarae* (in forest fragments) harbored the same *Borrelia* lineage (B3), challenging the ecological structuring observed for *Rickettsia*. Given that host sharing has been documented in the *Borrelia* system elsewhere (46), we hypothesize that this pattern reflects broad host susceptibility of the B3 lineage. Presence of such shared lineage along with other potentially pathogenic SFG *Rickettsia* and RFG *Borrelia* in the rodent community suggests occupational exposure risks for individuals working in forest-adjacent environments, such as plantation laborers and forest users, who may encounter infected vectors (47).

### 4.3. Limited coinfection indicates largely independent transmission pathways

Coinfections involving multiple pathogens are common in wildlife and can have profound consequences for host fitness (48). In many host–pathogen systems, coinfecting agents interact through immune-mediated mechanisms, resulting in either synergistic or antagonistic effects on disease outcomes (49). Motivated by these insights, we examined the occurrence of *Rickettsia, Borrelia*, and *Bartonella* coinfections in the host community. However, coinfections were infrequent and showed no specific patterns, suggesting limited interaction among these bacteria and independent vector–host transmission pathways. Moreover, the absence of sex-based or species-specific differences in infection probability implies that exposure risk is shaped by other factors, potentially including variation in vector contact and their seasonality, rather than by sex-specific behaviors (50). However, several important limitations should be considered when interpreting these results, including limited sample sizes for certain hosts (e.g., *R. rattus*) and the absence of ectoparasite screening for these focal bacterial genera. Additionally, reliance on single-gene phylogenetic inference alone cannot definitively establish the pathogenicity of the SFG and RFG lineages. Future studies integrating vector surveillance with experimental and pathological approaches will be essential to establish the clinical relevance of these bacterial lineages and to more accurately assess their potential risk to wildlife, livestock, and humans.

## 5. Conclusion

In this study, we characterize SFG *Rickettsia* and RFG *Borrelia* circulating in rodents of southern India. Although prevalence was low, the detection of genetically diverse lineages suggests diversity that is underrepresented and often overlooked in clinical studies. Distinct tissue associations, host specificity, and limited coinfection indicate that infection patterns are shaped by host ecology, pathogen biology, and vector interactions rather than uniform exposure across host species. By providing baseline data from wild and synanthropic rodents, this study highlights the value of ecologically informed wildlife surveillance to improve understanding of the etiology of undiagnosed infections and zoonotic risk.

## Acknowledgements

This study was supported by the Department of Atomic Energy, Government of India (Project Identification RTI 4006) awarded to UR. We are grateful to K.M. Cariappa of Kadamane for facilitating the fieldwork. We thank S. Vijayakumar and K. Kamaraj for their assistance with field sampling, and Dr. Sandhya Ganesan (Indian Institute of Science Education and Research Thiruvananthapuram) for providing the positive control for *Coxiella*. We also acknowledge the support of the NCBS Sanger Sequencing Facility.

## Ethics statement

This study was approved by the NCBS Institutional Animal Ethics Committee (NCBS-IAEC-2016/10-[M]). Sample collection and laboratory processing in this study adhered to strict biosafety guidelines and were approved by the Institutional Biosafety Committee (TFR: NCBS:23_IBSC/2017).

## Data availability

Sequence data generated in this study have been deposited in GenBank under the accession numbers PX803039-PX803047, PX788653-PX788660, and PX788662.

## Biography

Dr. Ansil is a postdoctoral fellow in the School of Biological Sciences at the University of Oklahoma. His work focuses on how environmental change influences zoonotic pathogen dynamics in wildlife, with a special interest in rodent and bat systems in the tropics.

