## Supplementary Tables for "Spotted fever *Rickettsia* and relapsing fever *Borrelia* in rodents from southern India"

**S1 Table.** Rodents sampled across habitat types within the Kadmanae landscape and screened for *Rickettsia*, *Borrelia*, *Orientia*, *Leptospira*, and *Coxiella*.

| Species | Forest | Grassland | Human habitation | Total |
|---|---|---|---|---|
| *Rattus rattus* | - | - | - | 10 |
| *Rattus satarae* | 66 | - | - | 66 |
| *Mus cf. fernandoni* | 2 | 20 | - | 22 |
| *Mus cf. famulus* | 2 | 19 | 5 | 26 |

**S2 Table.** Primers and PCR conditions used for different bacterial detection. *Borrelia* and *Leptospira* screening followed a nested PCR approach. '*' and '+' indicate outer and inner primer sets of nested protocol, respectively.

| Bacterial genera | Locus | Primer sequence (5'-3') | Size (bp) | Annealing temperature | Reference |
|---|---|---|---|---|---|
| *Rickettsia* | ompB | 120-2788F: AAACAATAATCAAGGTACTGT | 811 | 58°C | (1) |
| | | 120-3599R: TACTTCCGGTTACAGCAAAGT | | | |
| *Borrelia* | 16S rRNA | 16S1AF: CTAACGCTGGCAGTGCGTCTTAAGC* | 720 | 63°C | (2) |
| | | 16S1BR: AGCGTCAGTCTTGACCCAGAAGTTC* | | | |
| | | 16S2AF: AGTCAAACGGGATGTAGCAATAC+ | 650 | 61°C | |
| | | 16S2BR: GGTATTCTTTCTGATATCAACAG+ | | | |
| *Orientia* | 56-kDa protein | OtsuF: AATTGCTAGTGCAATGTCTG | 371 | 55°C | (3) |
| | | OtsuR: GGCAT- TATAGTAGGCTGAG | | | |
| *Leptosira* | 16S rRNA | LeptoAF: GGCGGCGCGTCTTAAACATG* | 330 | 63°C | (4) |
| | | LeptoBR: TTCCCCCCATTGAGCAAGATT* | | | |
| | | LeptoCF: CAAGTCAAGCGGAGTAGCAA+ | 289 | 63°C | |
| | | LeptoDR: CTTAACCTGCTGCCTCCCGTA+ | | | |
| *Coxiella* | IS1111 A | F: CGCAGCACGTCAAACCG | 146 | 60°C | (5) |
| | | R: TATCTTTAACAGCGCTTGAACGTC | | | |

**S3 Table:** Sample specific prevalence of *Rickettsia* spp, *Borrelia* spp, and *Leptospira* spp. Samples tested for *Coxiella* spp (spleen and pooled tissue) and *Orientia* spp (blood and pooled tissue) were negative.

| | | ***Rickettsia*** | | ***Borrelia*** | | ***Leptospira*** | |
|---|---|---|---|---|---|---|---|
| **Species** | **Sample type** | **% (pos/n)** | **95% CIs** | **% (pos/n)** | **95% CIs** | **% (pos/n)** | **95% CIs** |
| *Rattus rattus* | Blood | 0 (0/10) | 0- 27.75 | 10 (1/10) | 0.51- 40.41 | - | - |
| | Pooled tissue | 10 (1/10) | 0.51- 40.41 | 0 (0/10) | 0- 27.75 | 0 (0/10) | 0- 27.75 |
| | Kidney | - | - | - | - | 0 (0/10) | 0- 27.75 |
| *Rattus satarae* | Blood | 0 (0/66) | 0- 5.50 | 6.06 (4/66) | 2.38- 14.57 | - | - |
| | Pooled tissue | 12.12 (8/66) | 6.27- 22.13 | 0 (0/66) | 0- 5.50 | 1.51 (1/66) | 0.07- 8.09 |
| | Kidney | - | - | - | - | 1.51 (1/66) | 0.07- 8.09 |
| *Mus cf. fernandoni* | Blood | 0 (0/22) | 0- 14.86 | 13.63 (3/22) | 4.74- 33.33 | - | - |
| | Pooled tissue | 0 (0/22) | 0- 14.86 | 0 (0/22) | 0- 14.86 | 0 (0/22) | 0- 14.86 |
| | Kidney | - | - | - | - | 0 (0/22) | 0- 14.86 |
| *Mus cf. famulus* | Blood | 0 (0/26) | 0- 12.87 | 0 (0/26) | 0- 12.87 | - | - |
| | Pooled tissue | 0 (0/26) | 0- 12.87 | 0 (0/26) | 0- 12.87 | 0 (0/26) | 0- 12.87 |
| | Kidney | - | - | - | - | 0 (0/26) | 0- 12.87 |
| | **Total** | **7.26 (9/124)** | **3.86- 13.21** | **6.45 (8/124)** | **3.30- 12.21** | **0.80 (1/124)** | **0.04- 4.42** |

**S4 Table:** Set of generalized linear models (GLMs) used to evaluate the effects of species, sex, and their interaction on infection status for *Rickettsia* and *Borrelia*. Models were ranked using ΔAICc, with the number of parameters (*k*) and Akaike weights (*wi*) reported for each model.

| **Model** | *k* | ΔAICc | *wi* |
|---|---|---|---|
| Rickettsia infection status ~ species + sex | 5 | 0 | 0.967 |
| Rickettsia infection status ~ species * sex | 8 | 6.75 | 0.033 |
| Borrelia infection status ~ species + sex | 5 | 0 | 0.958 |
| Borrelia infection status ~ species * sex | 8 | 6.23 | 0.042 |
