## Supplementary Figures for "Spotted fever *Rickettsia* and relapsing fever *Borrelia* in rodents from southern India"

**Figure S1:** *Rickettsia, Borrelia, Leptospira* and *Bartonella* infection status among individual rodents. Bartonella infection status was collated from (1)

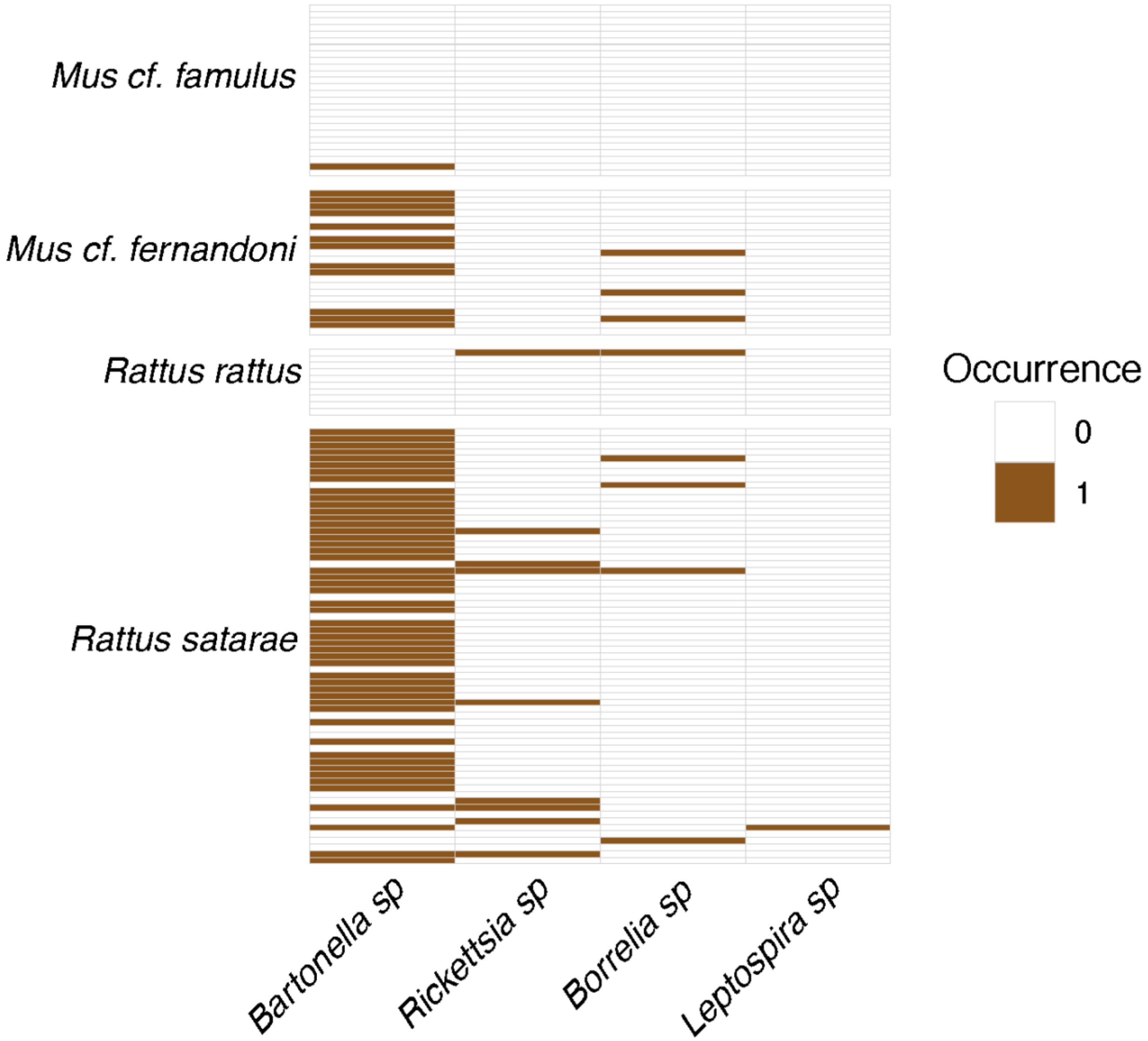

**Figure S2:** (A) Maximum-likelihood phylogenetic tree of *Leptospira* spp. based on 16S rRNA gene sequences, constructed using 10,000 bootstrap replicates under the GTR + G + I substitution model. (B) Genetic similarity of the *Leptospira* sequence derived from *Rattus satarae* in comparison with closely related *Leptospira* species.

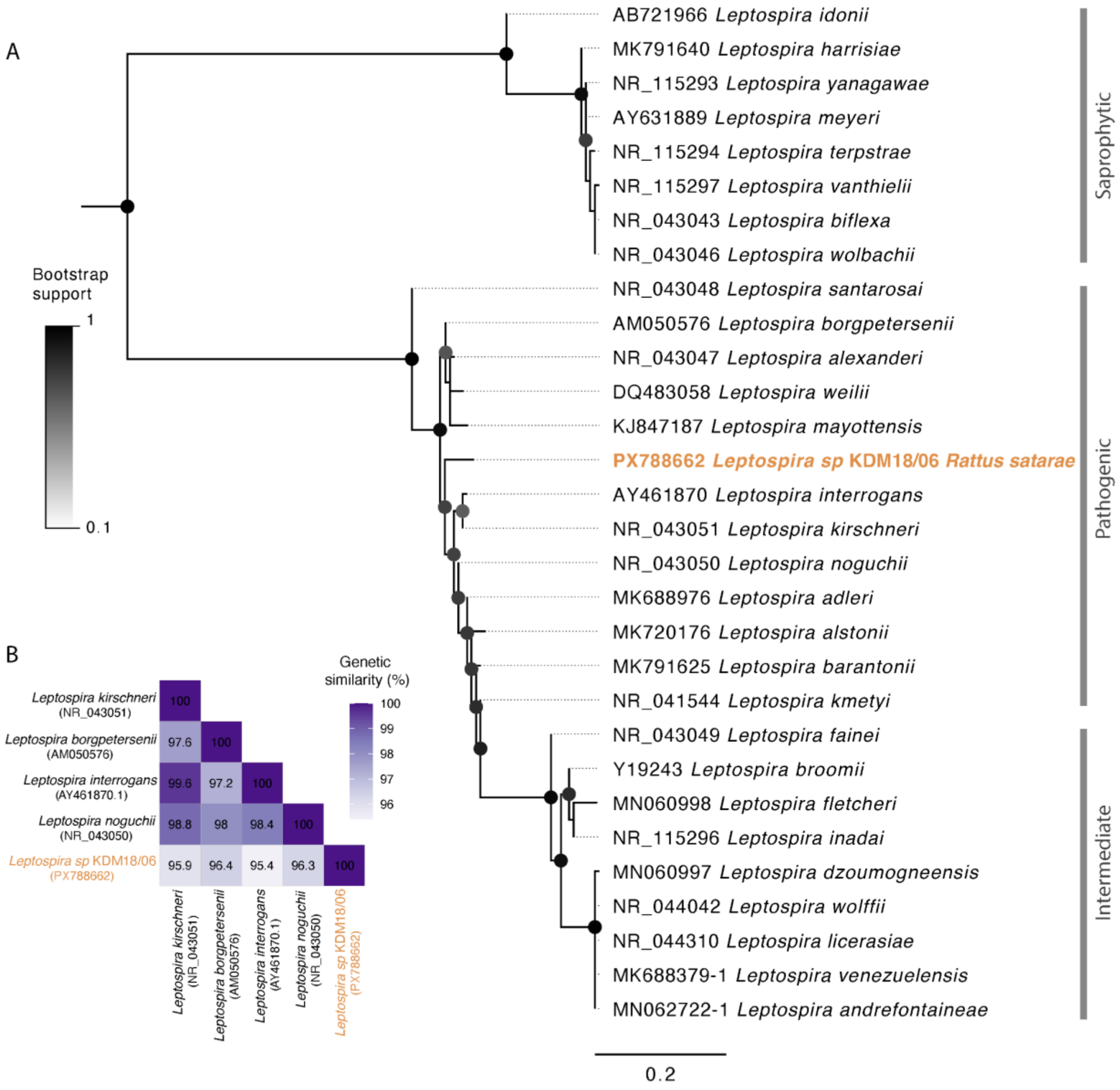
